# Chromosome segregation drives division site selection in *Streptococcus pneumoniae*

**DOI:** 10.1101/087627

**Authors:** Renske van Raaphorst, Morten Kjos, Jan-Willem Veening

## Abstract

Accurate spatial and temporal positioning of the tubulin-like protein FtsZ is key for proper bacterial cell division. *Streptococcus pneumoniae* (pneumococcus) is an oval-shaped, symmetrically dividing human pathogen lacking the canonical systems for division site control (nucleoid occlusion and the Min-system). Recently, the early division protein MapZ was identified and implicated in pneumococcal division site selection. We show that MapZ is important for proper division plane selection; thus the question remains what drives pneumococcal division site selection. By mapping the cell cycle in detail, we show that directly after replication both chromosomal origin regions localize to the future cell division sites, prior to FtsZ. Perturbing the longitudinal chromosomal organization by mutating the condensin SMC, by CRISPR/Cas9-mediated chromosome cutting or by poisoning DNA decatenation resulted in mistiming of MapZ and FtsZ positioning and subsequent cell elongation. Together, we demonstrate an intimate relationship between DNA replication, chromosome segregation and division site selection in the pneumococcus, providing a simple way to ensure equally sized daughter cells without the necessity for additional protein factors.

## Introduction

In eukaryotic cells, DNA replication, chromosome segregation and cell division are tightly coordinated and separated in time (*1*–*3*). In most bacteria this is less obvious as these processes occur simultaneously. However, in the last decade, it has become evident that the bacterial cell cycle is a highly regulated process, in which both cell cycle proteins as well as the chromosome have defined spatial and temporal localization patterns (*4*, *5*). The tubulin-like protein FtsZ (forming the Z-ring) is key for initiating divisome assembly in virtually all bacteria (*6*). Accurate cell division is mostly exerted through regulation of FtsZ positioning in the cell. However, the mechanisms that control FtsZ positioning can be highly diverse between bacterial species. In well-studied rod-shaped model organisms, such as *Bacillus subtilis* and *Escherichia coli*, precise formation of the Z-ring at midcell is regulated by the so-called Min-system and nucleoid occlusion (*7*, *8*). These are both negative regulators of FtsZ polymerization, which prevent premature Z-ring formation and cell division near cell poles and across unsegregated chromosomes, respectively. These two regulatory mechanisms have been found in many bacteria. However, in some species other dedicated proteins are used for this purpose, including MipZ in *Caulobacter crescentus* (*9*), SsgB in *Streptomyces coelicolor*(*10*) and PomZ in *Myxococcus xanthus* (*11*). It is important to note, however, that none of these FtsZ regulation mechanisms are essential for bacterial growth, and other mechanisms of cell cycle control must therefore also exist (*12*–*14*). In this context, is has been suggested that there are important links between different cell cycle processes, such as DNA replication and Z-ring assembly (*13*–*16*).

As for the opportunistic pathogen *S. pneumoniae,* the orchestration of replication and chromosome dynamics remains largely unknown. Ovococcal *S. pneumoniae* lack a nucleoid occlusion system and has no Min-system (*17*, *18*). Recently, the putative division site selector MapZ (or LocZ) was identified in *S. pneumoniae* (*19*, *20*). This protein localizes early at new cell division sites and positions FtsZ by a direct protein-protein interaction (*19*). MapZ is binding peptidoglycan (PG) via an extracellular domain, and is also a target of the master regulator of pneumococcal cell shape, the Ser/Thr kinase StkP (*19*–*21*). Together, this suggests that for division site selection in *S. pneumoniae*, FtsZ is controlled via the MapZ beacon at midcell (*13*, *22*, *23*). Furthermore, the mechanisms of chromosome segregation in pneumococci also seem to be different than in rod-shaped model bacteria; *S. pneumoniae* harbors a single circular chromosome with a partial partitioning system that only contains the DNA-binding protein ParB with *parS* binding sites, but lacks the ATPase ParA. Furthermore, the ubiquitous condensin protein SMC is not essential (*24*). Although both ParB and SMC are involved in chromosome segregation in pneumococci, *parB* and *smc* mutants have minor growth defects and a low percentage of anucleate cells (1-4 %) (*24*, *25*). In contrast, in *B. subtilis* deletion of *smc* is lethal at normal growth conditions (*26*). To gain more understanding of the progression of the pneumococcal cell cycle, we therefore investigated the relationship between DNA replication, chromosome segregation and division site selection in the pneumococcus. We show that MapZ is not involved in division site selection as suggested before, but is crucial for correctly placing the Z-ring perpendicularly to the length axis of the cell. By establishing new tools to visualize the replisome and different genetic loci, we show that there is an intimate relationship between DNA replication, chromosome segregation and division. Importantly, we demonstrate that correct chromosomal organization acts as a roadmap for accurate division site selection in pneumococcus and possibly other bacteria.

## Results

### MapZ sets the pneumococcal division plane

In contrast to what can be expected for a protein involved in division site selection, *ΔmapZ* mutants are not elongated but on average shorter than wild type cells (*19*, *20*) with relatively minor distortions in cell morphology (*20*, *27*), raising questions on what the actual biological function of MapZ is (*27*). To reassess the *ΔmapZ* phenotype, we fused MapZ at its N-terminus to a monomeric superfolder green fluorescent protein (GFP). Using the cell-segmentation software Oufti (*28*), to detect cell outlines and fluorescent signals, in combination with the newly developed R-package SpotprocessR to analyze the microscopy data (see Methods), GFP-MapZ localization was determined in exponentially growing cells (balanced growth). Note that balanced growth, by re-diluting exponentially growing cells several times, pneumococcal cell length becomes an accurate proxy for the cell cycle state (*18*, *21*). Cells were ordered by length, and this order was plotted as a density plot against the position of GFP-MapZ on the long axis of the cells (Fig. 1A). In line with previous reports (*19*, *20*), GFP-MapZ localized to the division site (Fig. 1A). As new cell wall is synthesized at midcell (*17*), MapZ seems to move along with the current division site, probably via attachment to PG, and ends up at the interface between the new and old cell halves. This position will eventually become the future division site where the Z-ring assembles. Deleting *mapZ* in the encapsulated D39 genetic background led to irregularly shaped and shorter, sometimes branched or clustered cells (Fig. 1B). Similar observations were made in serotype 4 strain TIGR4 and in the unencapsulated R6 laboratory strain (fig. S1A and B). To examine FtsZ localization, we constructed a C-terminal monomeric red fluorescent protein (mCherry) fusion to FtsZ expressed from its own locus as the only copy (*29*). While the FtsZ-mCherry strain showed a normal cell size distribution in a wild type background, when combined with the *ΔmapZ* mutant, a clear synthetic phenotype arose and cells were misformed (fig. S2A and B), suggesting that the previously described *mapZ* phenotype in the presence of FtsZ-fusions should be interpreted with caution (*27*). Therefore, we replaced FtsZ-mCherry by a functional FtsZ-CFP or FtsZ-mKate2 (FtsZ-RFP) fusion (fig. S2C and D), and reassessed FtsZ localization in *ΔmapZ* cells. As reported before (*19*, *20*), localization of FtsZ to future division sites occurs when MapZ is already localized at this position (Fig. 1A and C). Note, however, that in stark contrast to MapZ, which gradually moves as new cell wall is synthesized, FtsZ is highly dynamic and remodels quickly from the previous to the future division site. Thus, there is only a brief moment in the cell cycle where MapZ and FtsZ colocalize (cf. Fig. 1A with Fig.1C). Importantly, FtsZ localization over the length axis of the cell was not affected in *ΔmapZ* cells, suggesting that MapZ is not essential for accurate timing of Z-ring assembly. To gain more insights into the role of MapZ during septum formation, we stained cells with fluorescently-labeled vancomycin (Van-FL) to image sites of cell wall synthesis (*30*). By measuring the angle of the areas in the cell enriched with Van-FL relative to the long axis, we observed that the septum was formed perpendicular to the cell axis in the wild type (median deviation from 90 degrees = 3.08, se = 1.47 degrees), while in *ΔmapZ* cells this angle was skewed (Fig. 1D, median deviation from 90 degrees = 7.65, se = 1.27 degrees, significant difference p = 0.014, Kolmogorov-Smirnov test). Measuring the angle of FtsZ-CFP in the same manner confirmed that the angle of the Z-ring was skewed in *ΔmapZ* cells (fig. S2E). These results are in line with previous observations (*19*, *20*) and could explain the variability in cell shapes observed in *ΔmapZ* mutants. The observed cell shape alterations are reminiscent of *E. coli* mutants lacking certain low molecular weight penicillin binding proteins (LMW-PBPs) such as PBP5 that have defected division plane selection and mislocalized Z-rings (*31*). LMW-PBPs modify PG by trimming amino acid linkages from the glycan side chains (*32*). Since MapZ has a large extracellular PG binding domain and is controlled by the Ser/Thr kinase StkP (*19*, *33*), which is proposed to be a key player in tuning peripheral and septal peptidoglycan synthesis (*21*, *34*), it is tempting to speculate that MapZ has a function in cell wall remodeling and subsequently maintaining the perpendicular Z-ring plane.

**Fig. 1.**
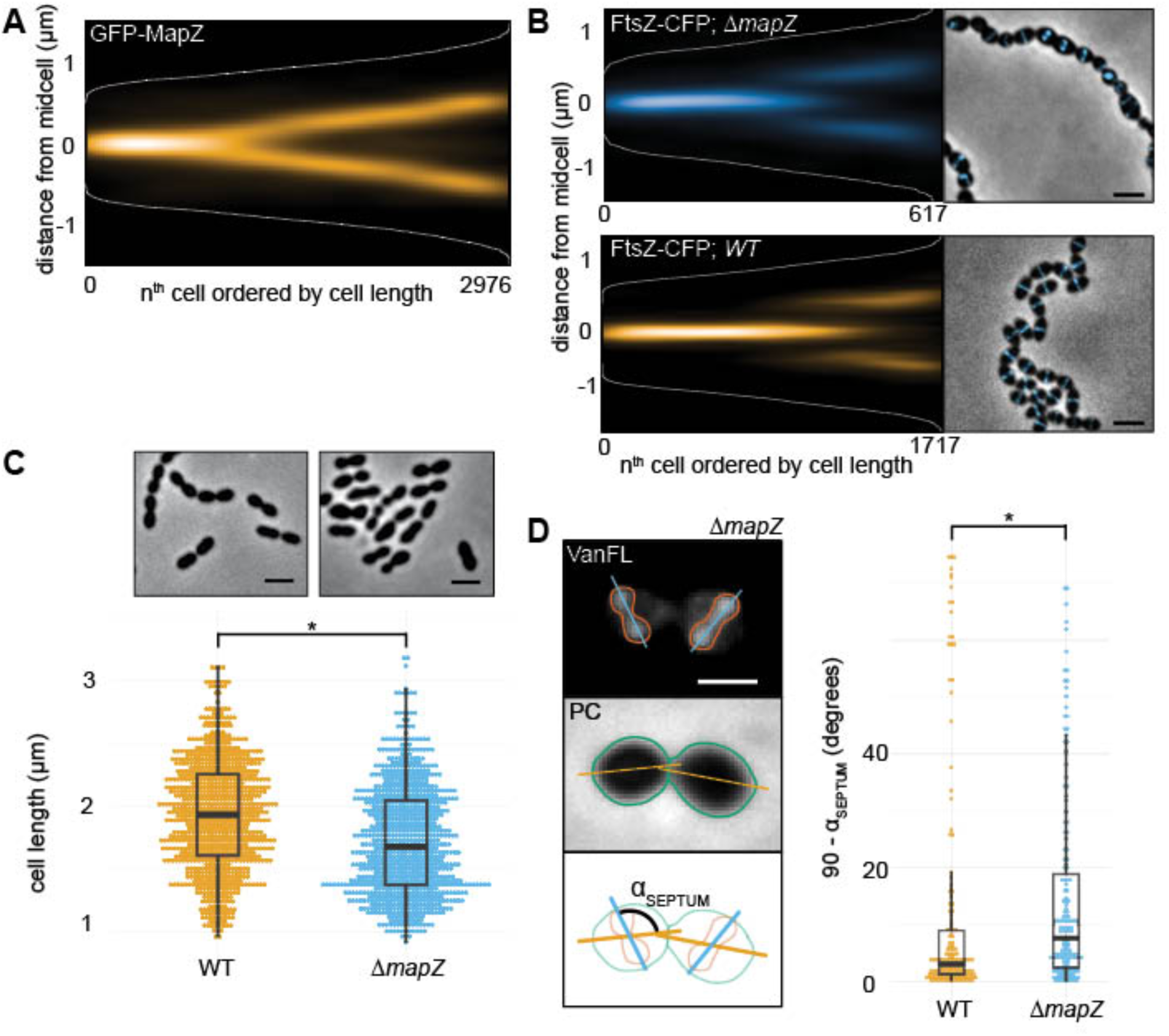
MapZ sets the pneumococcal division plane but is not involved in division site selection. (**A**) Fluorescence microscopy of 2976 cells, 7020 spot localizations were quantified and analyzed using Ouftie & SpotprocessR (see Methods). The distance of GFP-MapZ (strain RR101) from midcell was plotted in a heatmap where all localizations are ordered by cell length, and the color saturation represents protein density. MapZ is present at midcell at an early stage during the cell cycle. (**B**) Cell size distribution of wild type D39 and *ΔmapZ* cells (strain RR93), respresenting measurements of 1692 and 705 cells, respectively. Top: phase contrast microscopy images. Scale bar is 2 μm. (**C).** The localization of FtsZ-CFP in wild type (strain RR23) and *ΔmapZ* (strain RR105) cells as shown by histograms and micrographs from overlays of phase contrast images with CFP signal. Scale bar is 2 μm. The plots are based on data from 617 cells/957 localizations for *ΔmapZ* and 1717 cells/2328 localizations for wild type. (**D).** The angle of the septum relative to the length axis of the cells is less precise in *ΔmapZ* cells. Left: wild type D39 and *ΔmapZ* cells (strain RR93) were stained with fluorescently labeled vancomycin (VanFL). Fluorescence image (top), phase contrast image (middle) and a schematic drawing of the analysis (bottom) are shown. The angle, α_SEPTUM_, was measured automatically by measuring the angle between the long axis of the bounding boxes of the cell outlines and the long axis of the bounding box of the VanFL signal. Scale bar is 1 μm. Right: the α_SEPTUM_ plotted in wild type cells and *∆mapZ* cells. 177 and 181 cells were measured for wild type and *ΔmapZ*, respectively.

### The replisome of *S. pneumoniae* is dynamic around midcell

Since the *ΔmapZ* mutant has moderate effects on division site selection under our experimental conditions, another system must be in place. Since *S. pneumoniae* lacks the canonical systems, we hypothesized that ovococci might coordinate division via chromosome replication and segregation (*15*). To test this, we first aimed at imaging the DNA replication machine (replisome) and constructed inducible, ectopic fusions of the single-strand binding protein (SSB), the β sliding clamp (DnaN) and the clamp loader (DnaX) with GFP or RFP (mKate2). Fluorescence microscopy showed enriched signals as bright diffraction limited spots for all fusions, mainly localized in the middle of the cells, similar to what has been observed for *B. subtilis* and *E. coli* (*35*, *36*) (Fig. 2A). Notably, the background signal of SSB-RFP was higher than the background of the other fusions, as also reported for *E. coli* (*36*). Chromosomal replacements of the fusion constructs with the original gene could only be obtained for *dnaX*, but not for *ssb* and *dnaN*, suggesting that the fusion tags of these two latter genes are not fully functional. To validate that the localizations of the fusions represent biologically active replisomes, we examined their colocalization patterns. As expected, the ectopically expressed fluorescent fusions of DnaX, DnaN and SSB to RFP colocalize with the functional DnaX-GFP fusion in live cells (91 % colocalization or more, fig. S3A).

**Fig. 2.**
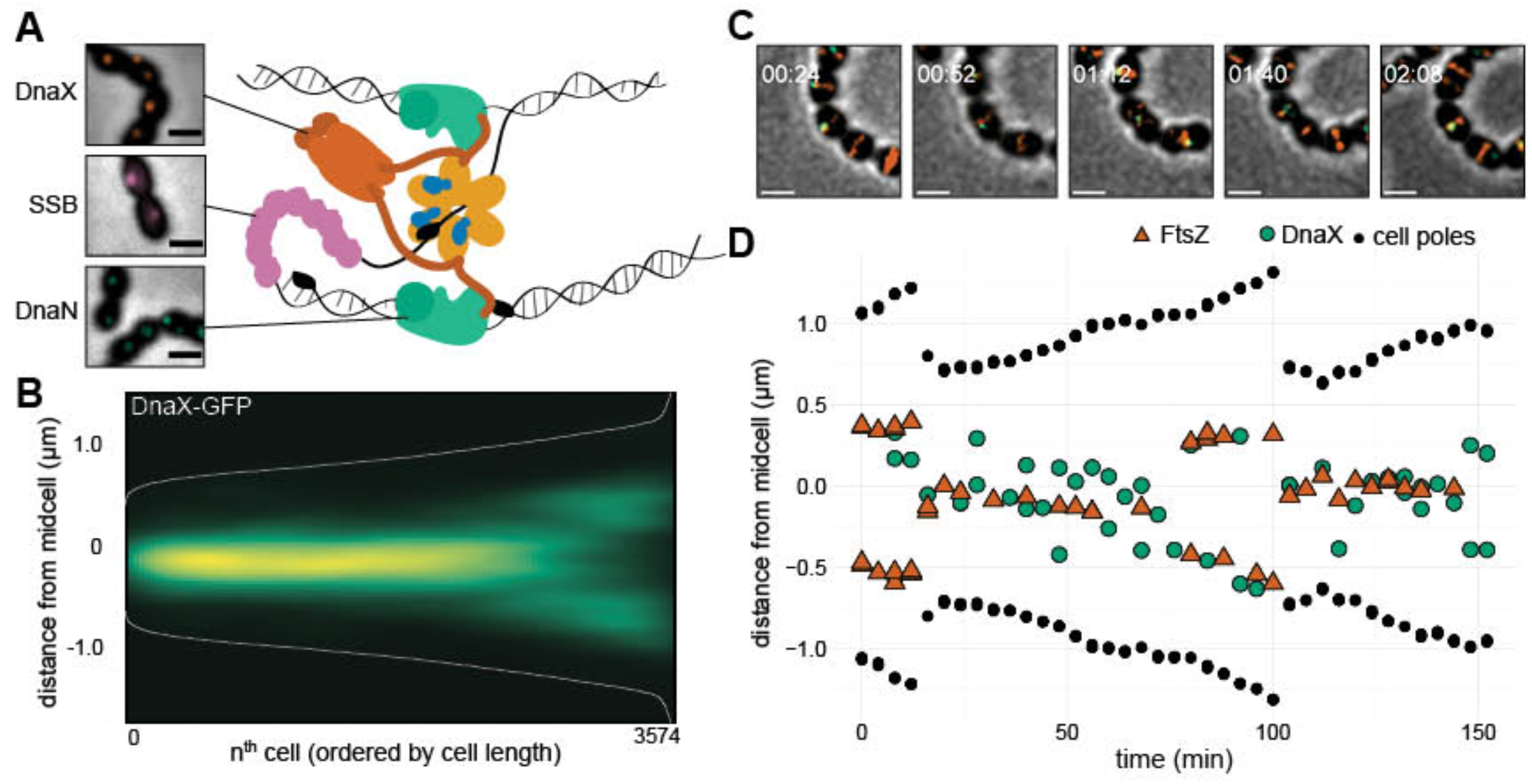
Localization of the pneumococcal replisome. (**A**) Localization of DnaX-GFP (RR31), GFP-DnaN (DJS02) and SSB-GFP (RR33). A cartoon of the bacterial replication fork shows the role of DnaX (clamp loader), SSB (single-strand binding protein) and DnaN (β sliding clamp). (**B**) Plotting the localization of DnaX-GFP (RR22) shows that the replisome is localized at midcell. Data from a total of 3574 cells, 3214 unique localizations. (**C**) Snap shots from a representative time-lapse movie of strain MK396 (*dnaX-GFP, ftsZ***-***mKate2*). Overlays of GFP, RFP and phase contrast are shown. Scale bar is 1 μm (**D**) Transcript of the cell shown in Fig. 2C. The distance of FtsZ (red), DnaX (green) and the cell poles (black) to midcell is plotted against time. The data is also shown in Movie S1. Transcripts of more single cells are shown in fig. S3.

In order to study the localization and dynamics of DnaX-GFP relative to the cell poles and the Z-ring, a DnaX-GFP/FtsZ-mKate2 double-labeled strain was made and exponentially growing cells were analyzed by fluorescence microscopy. These plots show that DnaX-GFP is positioned close to midcell with a similar pattern as FtsZ-mKate2, although the DnaX-GFP localization is less precise than FtsZ-RFP (Fig. 2B). To validate these results, tracking single cells during growth using time-lapse fluorescence microscopy (Fig. 2D and D, fig. S3B, Movie S1) showed that although the replisome(s) is dynamic, it stays in near proximity to the Z-ring. Surprisingly, the replisomes move to the future division sites with the same timing as FtsZ, and the Z-ring does not linger for cell division to finish (Fig. 2D).

To gain more insight into the nature of the movement of the replisome, we imaged DnaX-GFP in short-time interval movies (1 sec, Movie S2) using total internal reflection fluorescence (TIRF) microscopy. We tracked DnaX-GFP using the single molecule tracking software U-track (*37*) and analyzed mobility using SpotprocessR (fig. S3C). As expected, replisome mobility was significantly lower than that of free diffusing GFP (*38*). However, compared to ParB-GFP, which binds to the origin of replication (*oriC*) (*24*), DnaX-GFP showed a nearly two-fold higher mobility (MSD = 2.66 *10^-2^ μm^2^, D_*app*_ = 2.44*10^-3^μm^2^/s^-1^ compared to MSD = 8.8 * 10^-3^ μm^2^, D_*app*_ = 3.19*10^-4^ μm^2^/s^-1^; fig. S3D and D), indicating that DnaX movement is not strictly confined by the chromosome.

### The pneumococcal chromosome segregates in a longitudinal fashion

The on average midcell localization of the replication machineries in *S. pneumoniae* suggests that DNA replication at midcell might determine an ordered chromosomal organization. To examine this, methods for tagging chromosome positions in this bacterium were established (figs. S4 and S5). We first constructed a novel chromosome marker system based on fluorescent protein fusions to ParB of plasmid pLP712 (*39*) from *Lactococcus lactis* (hereafter named ParB_p_), which was found to require insertion of only a 18 bp *parS* binding site (hereafter named *parS*_*p*_) in the pneumococcal genome for visualizing genetic loci by microscopy (figs. S4A-C). The *parS*_*p*_ sequence is simpler compared to existing ParB/*parS* systems and does not require additional host factors (*40*, *41*). Importantly, our system does not perturb DNA replication and is completely orthogonal to *S. pneumoniae* chromosomal ParB (figs. S4B and S5A-D). Secondly, we adapted the TetR/*tetO* fluorescence repressor-operator system (FROS) (*42*) for *S. pneumoniae* and validated that it does not interfere with DNA replication (figs. S4D and S5A-C). To verify the localization patterns and compatibility of both systems, we constructed a strain containing both *parS*_*p*_ and *tetO* near *oriC* and showed that *parB*_*p*_-*gfp* and *tetR-rfp* foci colocalize (figs. S5E and F).

In total, five chromosomal locations were marked using ParB_p_/*parS*_*p*_ and/or TetR/*tetO*; the origin-region (359° degrees), right arm (101°), *ter*-region (178° and 182°) and two positions on the left arm (259° and 295°) (Fig. 3A). Using double-labeled strains under balanced growth, the localizations of loci were compared revealing that the pneumococcal genome is organized in a longitudinal fashion (Fig. 3B, fig. S6). The left and right arms move at the same time to the new daughter cells (Fig. 3B). The terminus region seemed less confined in space during the cell cycle (Fig. 3B). Strikingly, the origins never localized near the cell poles as is common in other bacteria (*5*, *42*–*44*), and arrived to future division sites at a very early stage, before DnaX and FtsZ, with a similar timing as MapZ (Fig 3B, left panel, D and E). The early segregation of *oriC* was also observed when single cells were tracked over time (Fig. 3F, fig. S7 and Movie S3).

**Fig. 3.**
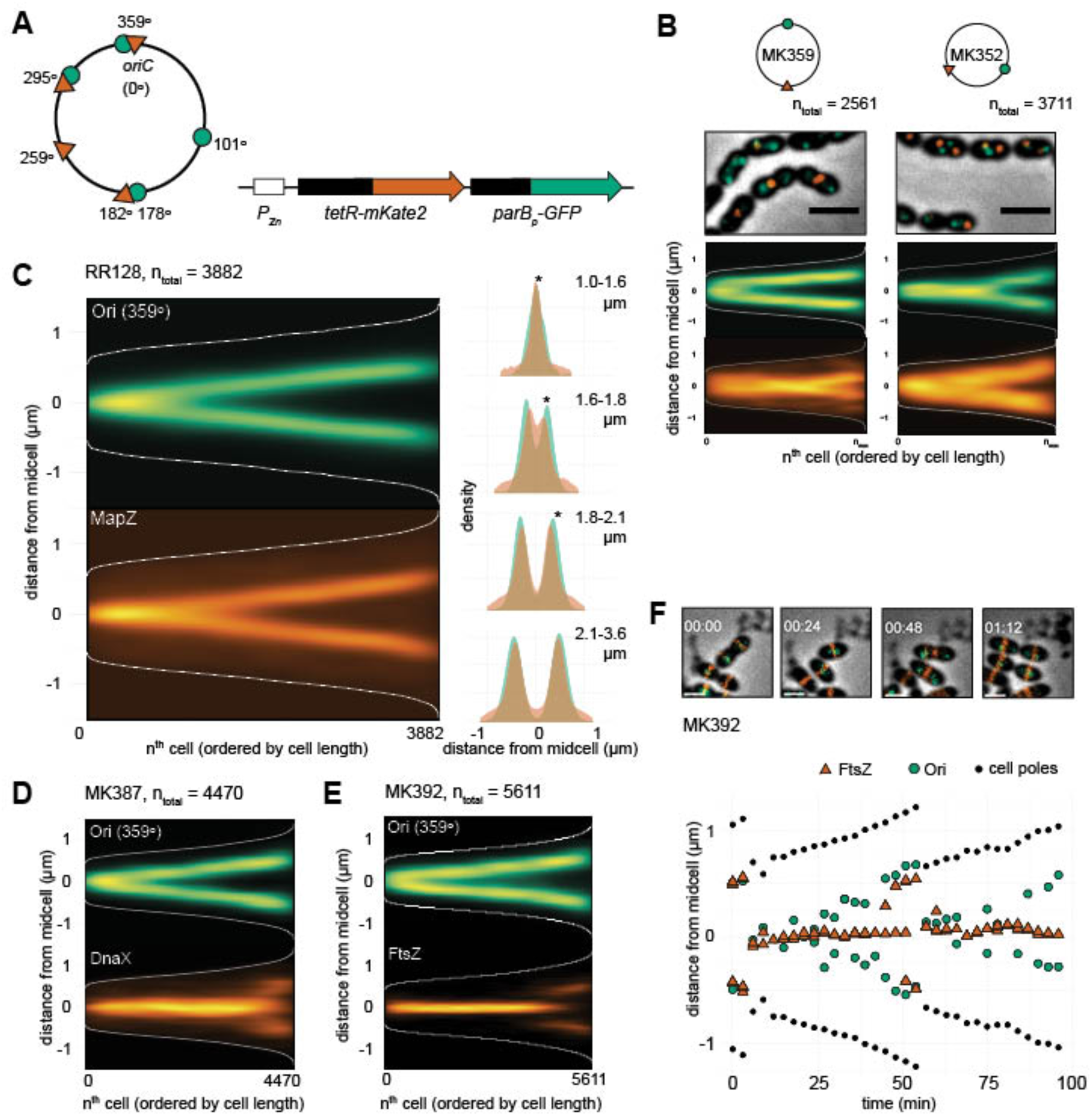
Chromosomal organization in *S. pneumoniae*. **(A)** Visualizing specific genetic loci in live cells by fluorescence microscopy was done by developing two independent chromosomal markers systems; TetR-mKate/*tetO* (*tetO* integration sites indicated by red triangles on the chromosome map) and ParB_p_-GFP/*parS*_*p*_ (*parS*_*p*_ integration sites indicated by green circles). *tetR-RFP* and *parB*_*p*_-*xGFP* are ectopically expressed from the non-essential *bgaA* locus under control of the Zn^2+^-inducible promoter P_Zn_. (**B**) Localization of the origin and terminus (MK359, left panel) and left and right arm (MK352, right panel) in exponentially growing cells. Overlays of GFP signal, RFP signals and phase contrast images are shown. Scale bars are 2 μm. The data represents 2561 cells/3815 GFP-localizations/2793 RFP-localizations from MK359 and 3711 cells/5288 GFP-localizations/4372 RFP-localizations from MK352. (**C**) Localization of the origin (ParB_p_-GFP/*parS*_*p*_ at 359°) and RFP-MapZ (RR128) on the length-axis of the cell shown as heatmaps (left) and overlay of both localization density plots when the cells are grouped in four quartiles by cell length (right). Stars indicate a significant difference between GFP and RFP localization (Kolmogorov-Smirnov test, p<0.05). The data represents 3882 cells/1785 GFP-localizations/8984 RFP localizations. (**D**) Localization of the origin (ParB_p_-GFP/*parS*_*p*_ at 359°) and DnaX-RFP (MK387). The data represents 4470 cells/5877 GFP-localizations/4967 RFP-localizations. (**E**) Localization of the origin (ParB_p_-GFP/*parS*_p_ at 359°) and FtsZ-RFP (MK392). The data represents a total of 5611 cells/6628 GFP-localizations/26674 RFP-localizations. (**F**) Time-lapse microscopy shows that the origins move to the next cell halves before FtsZ. Snap shots from a representative time-lapse experiment (top) and plotting of the distances of FtsZ, the origins and the cell poles relative to midcell in a single cell (bottom) are shown. More examples of origin and FtsZ localizations in single cells are shown in fig. S7.

### SMC is required for correct segregation of *oriC* and cell shape

The observation that *oriC* arrives at future cell division sites prior to FtsZ opens the question whether MapZ has a role in directing the chromosome. However, the origin still localized to the future division site in *ΔmapZ* (Fig. 4A). The localizations of MapZ and *oriC* were further analyzed in a wild type background by sorting the cells into four subgroups according to cell size and plotting the localizations as histograms over the cell lengths (Fig. 3C, right panel). This shows that MapZ is localized slightly closer to the old mid-cell in smaller, newborn cells (Fig. 3C, right panel, stars indicate a significant difference in the first three groups of cells, Kolmogorov-Smirnov test, p < 0.05), and that *oriC* localizes to the new mid-cell before MapZ. Given the early movement of the origin to the future cell division sites, we wondered whether instead the chromosome or nucleoid associated proteins could play a role in guiding Z-ring positioning.

**Fig. 4.**
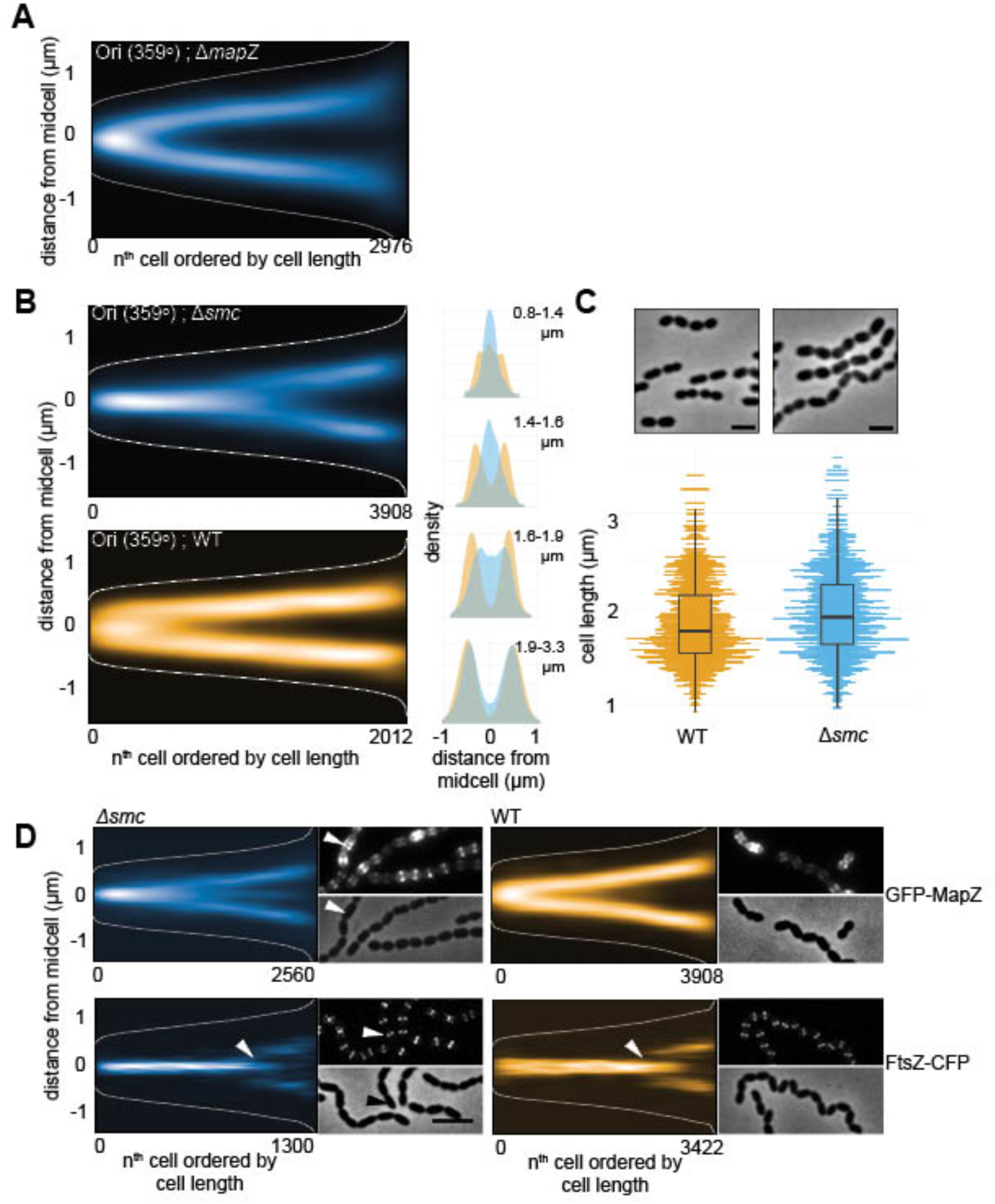
SMC is required for origin segregation and accurate division site selection. (**A**) Localization of the chromosomal origin (ParB_p_-GFP/*parS*_*p*_ at 359°) in a *ΔmapZ* background shows that MapZ does not affect origin segregation (RR99). The data represents 2976 cells/7020 localizations. (**B**) The origins (ParB_p_-GFP/*parS*_*p*_ at 359°) are segregated at a later stage in the cell cycle in *smc* compared to wild type. The localizations are shown as heatmaps when cells are sorted according to length (left) and as overlay of both localization density plots when the cells are grouped in four quartiles by cell length (right). The data represents 2012 cells/3815 localizations for wild type (MK359) and 3908 cells/5192 localizations for the *smc* mutant (MK368). (**C**) Phase contrast images of wild type D39 and *Δsmc* cells (AM39). The scale bar is 2 μm (top). Comparison of cell lengths between the wild type (1407 cell analyzed) and *Δsmc* (1035 cells analyzed, bottom). (**D**) Localization of GFP-MapZ and FtsZ-CFP in wild type versus *smc*. Fluorescence and phase contrast micrographs are shown along with heatmaps. The arrowhead in the micrograph points to a cell with clearly mislocalized MapZ. The arrowheads in the heatmap point the time when FtsZ remodels and assembles at the new division site in the wild type and in *Δsmc*. Data represents 2560 cells/5314 localizations (Δ*smc*, GFP-MapZ, RR110), 1300 cells/2257 localizations (Δ*smc*, FtsZ-CFP, RR84), 3908 cells/3128 localizations (wild type, GFP-MapZ, RR101) and 3422 cells/29464 localizations (wild type, FtsZ-CFP, RR70).

In many prokaryotes, condensin-like proteins called Structural Maintenance of Chromosomes (SMC) play a role in the organization and compaction of the chromosome (*45*). In *S. pneumoniae*, deletion of *smc* leads to approximately 2% anucleate cells and problems in chromosome segregation (*24*, *25*). In order to specifically investigate how the absence of SMC affected chromosome organization, the origin, terminus and left/right arm chromosome positions were determined in Δ*smc* cells. In line with what has been found using temperature-sensitive or degradable alleles in *B. subtilis* (*46*, *47*), the origin region arrived to the new midcell at a considerable delay when SMC was absent (Fig. 4B). Quantitative analysis of the origin localization showed a significant different localization of *oriC* in Δ*smc* vs wild type (p<1.5*10^-3^, Kolmogorov-Smirnov test, Fig. 4B). Eventually, however, the origins still segregated to their correct localization in subsequent larger cells. Also note that, segregation of the left arm, right arm and terminus did not differ significantly from wild type (fig. S8). Thus, *S. pneumoniae* SMC is specifically important for the early segregation of *oriC*.

In the current data, we also found that Δ*smc* cells are longer, more irregularly shaped and form long chains (Fig. 4B). The same observation was also made upon deletion of *smc* in strains TIGR4 and R6 (fig. S1A and B). This suggests that *smc* mutants are somehow defective, not only in chromosome segregation, but also in cell division. We therefore compared the localization of MapZ and FtsZ in wild type and Δ*smc* cells and found that the timing and accuracy of MapZ and FtsZ localization was altered in Δ*smc* (Fig. 4D). MapZ showed an obvious mislocalization; part of the MapZ-rings arrived at the new septa at a later stage, while a large fraction stayed at midcell in larger cells (11% of the MapZ-localizations in the largest quartile of the cell population stayed at midcell in ∆*smc* cells vs. 4% in wild type cells, Fig. 4d). For FtsZ, the effect is less pronounced, but a difference in localization accuracy is observed at the time when new Z-rings were formed (Fig. 4d). Note that the angle of the division ring is not affected in Δ*smc* cells (fig. S1C). Together these results suggest that SMC and/or origin localization is important for timely and precise positioning of the cell division machinery in *S. pneumoniae*.

Knowing that both origins and MapZ localize very early in the pneumococcal cell cycle to the future division sites and that perturbation of both of these individually cause division problems, we deleted both *smc* and *mapZ* in order to understand more about the link between them. The double mutant strain was readily obtained, although the strain had severe defects in growth and cell shape (fig. S1C and D). Notably, the phenotype of the double mutant looked like a combination of the single mutants; like *ΔmapZ*, the cells were on average smaller with large cell shape variation due to non-perpendicular division ring formation (fig. S1D), and like Δ*smc*, they displayed a chaining phenotype probably reflecting problems with timing of division ring formation leading to consecutive problems in timing of division and cell wall splitting (fig. S1D). These observations suggest that MapZ and SMC have independent roles in pneumococcal cell division, where SMC in important for timely localization of the division site, while MapZ is involved in correct, perpendicular placement of the division ring.

### Proper localization of *oriC* is crucial for division site selection in *S. pneumoniae*

We show above that deletion of *smc* caused a cell division defect in *S. pneumoniae* distinct from the *ΔmapZ* phenotype. To untangle whether this was a direct effect of SMC or whether it was caused by the resulting chromosome organization defect, we exposed *S. pneumoniae* to sublethal concentrations of ciprofloxacin in order to disturb chromosome organization while keeping *smc* intact. Ciprofloxacin is a broad-spectrum antibiotic which blocks the activities of type II topoisomerases and thereby affects DNA supercoiling and chromosome decatenation (*48*). Strikingly, when exponentially growing cells are transferred to a non-lethal concentration of ciprofloxacin (0.4 μg/mL), cells rapidly increase in cell length and form longer chains when compared to untreated cells (Fig. 5A and B). Origin splitting was clearly delayed in ciprofloxacin treated cells, and the timing and accuracy of Z-ring formation was severely affected (Fig. 5C). Moreover, localization of the replisome was less confined to the center of the cells, as was observed for *Δsmc* cells (fig. S9). Note that, at the ciprofloxacin concentration used in this experiment, replication elongation is reduced, but new rounds of replication are still initiated (*49*).

**Fig. 5.**
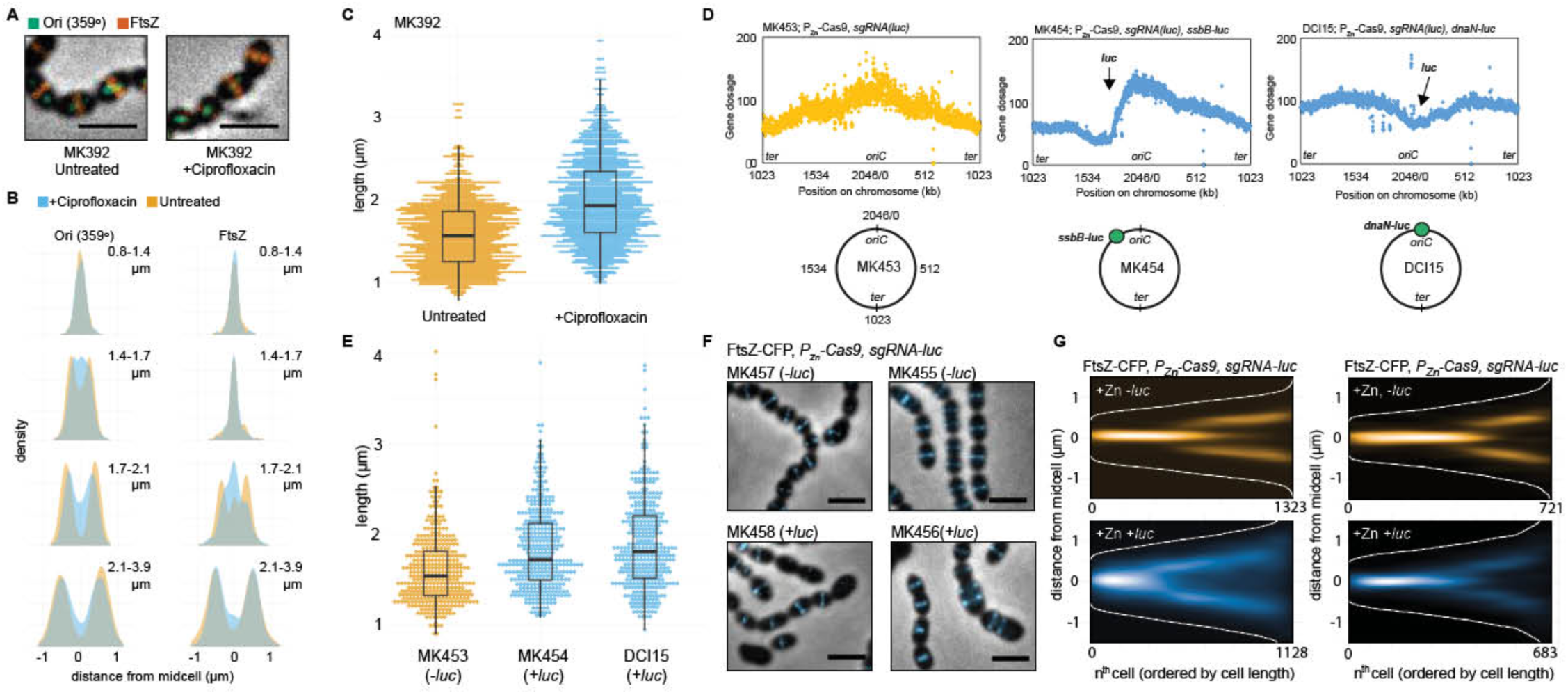
Perturbed chromosome segregation delays cell division. (**A-C**) Comparison of *S. pneumoniae* D39 wild type cells treated or untreated with sublethal concentrations (0.4 μg/ml) of ciprofloxacin for 60 min. (**A**) Images of strain MK392 (ParB_p_-GFP/*parS*_*p*_ at 359°, FtsZ-RFP) with overlay of phase contrast, GFP signals representing the origin and RFP signal representing FtsZ-RFP. Scale bar is 2 μm. (**B**) Subcellular localization of the origin (left) and FtsZ (right) in MK392 cells treated (blue) or untreated (yellow) with ciprofloxacin. Localization density plots when the cells are grouped in four quartiles by cell lengths are shown. Data represents 1138 cells/2518 RFP-localizations/2762 GFP-localizations for cells treated with ciprofloxacin and 1402 cells/2940 RFP-localizations/2540 GFP-localizations for untreated cells. (**C**) Cell length comparison of ciprofloxacin-treated (total 1138) and non-treated (total 1402) cells (MK392). (**D**-**G**). Comparisons of cells with or without Cas9-nuclease cut chromosome. The expression of Cas9 (together with a constitutively expressed single-guide RNA directed to the *luc*-gene) was induced in cells with or without the *luc* gene located on the chromosome. The *luc* gene was inserted either in the origin region (0°) or at the left arm (301°). (**D**) Whole genome marker frequency analysis of strains without *luc* (MK453), *luc* at 301° (MK454) or *luc* at the origin (DCI15). The number of mapped reads (gene dosage) is plotted as a function of the position on the circular chromosome. The chromosomal position of the inserted *luc* gene is indicated in the plot and on the schematic chromosome maps. (**E**) Cell size comparison of cells with and without cut chromosomes. The number of cells measured were 643 for the non-cut strain (MK453), 393 for the strain cut at 301° (MK454) and 383 for the strain cut near the origin (DCI15). (**F**) Overlay of FtsZ-CFP signals with phase contrast images show that cell morphologies are affected in cells with cut chromosomes. Scale bar is 2 μm. (**G**) Localization of FtsZ-CFP shown as heat maps where cells are ordered according to cell length. The data represents 1323 cells/3117 localizations for MK455, 1128 cells/3509 localizations for MK456, 721 cells/1133 localizations for MK457 and 683 cells/1209 localizations for MK458.

Finally, we also perturbed the DNA biology by cutting the chromosome at two different locations; close to *oriC* (at 0°) and on the left arm (at 295°), using an inducible CRISPR/Cas9 system (see Methods). Whole-genome marker frequency analysis of these strains after induction of Cas9, showed the expected cleavage of the chromosomal DNA at these two positions in the respective strains and major alterations in the replication patterns were observed (Fig. 5D). Cutting of the chromosome also caused severe problems with mistimed localization of FtsZ (Fig. 5E and F) and increased cell sizes (Fig. 5G). Interestingly, the effects on DNA replication were more pronounced when Cas9 was targeted to *oriC* compared to the left arm location, and, subsequently, proper control of Z-ring formation was completely lost in the former case. Together, these results further confirm that normal chromosome segregation, and origin segregation in particular, is key for well-timed Z-ring assembly and cell division progression in *S. pneumoniae*.

## Discussion

By detailed mapping of DNA replication and chromosome segregation in live *S. pneumoniae* cells, we found that proper segregation of the chromosomal origin is crucial for division site selection in this bacterium. We show that the pneumococcal chromosome is organized in a longitudinal fashion (ori-left/right-ter-ter-left/right-ori; Fig. 3 and fig. S6) with specific subcellular addresses for each locus. In contrast to *B. subtilis*, *C. crescentus* and fast-growing *E. coli*(*5*, *42*–*44*), the origins never localize near the cell poles in*S. pneumoniae*, and the organization is in this aspect more similar to the situation in slow-growing *E. coli* (*50*). Importantly, the newly replicated origins immediately mark the future cell division sites while the terminus remains at midcell. This organization is somewhat reminiscent of the chromosomal organization in *B. subtilis* and *E. coli* but is slightly simpler as every replicated locus eventually segregates to midcell before a new round of replication initiates (Fig. 6). Segregation of the chromosomal origin was directed by SMC and deletion of *smc* caused a marked delay in origin segregation, which in turn led to alterations in the timing of localization of important cell division proteins such as MapZ and FtsZ. Mistimed MapZ and FtsZ ultimately resulted in chain-forming cells with increased size. Importantly, the observed cell division defects are not caused by the deletion of *smc per se;* treatment of the cells with sublethal concentrations of ciprofloxacin or a CRISPR/Cas9-induced segregation block also caused similar cell division defects (ie. larger cells and chaining). Together, this indicates that timely segregation and positioning of the chromosomal origin at the quarter position in cells is important for orchestrating pneumococcal cell division.

**Fig. 6.**
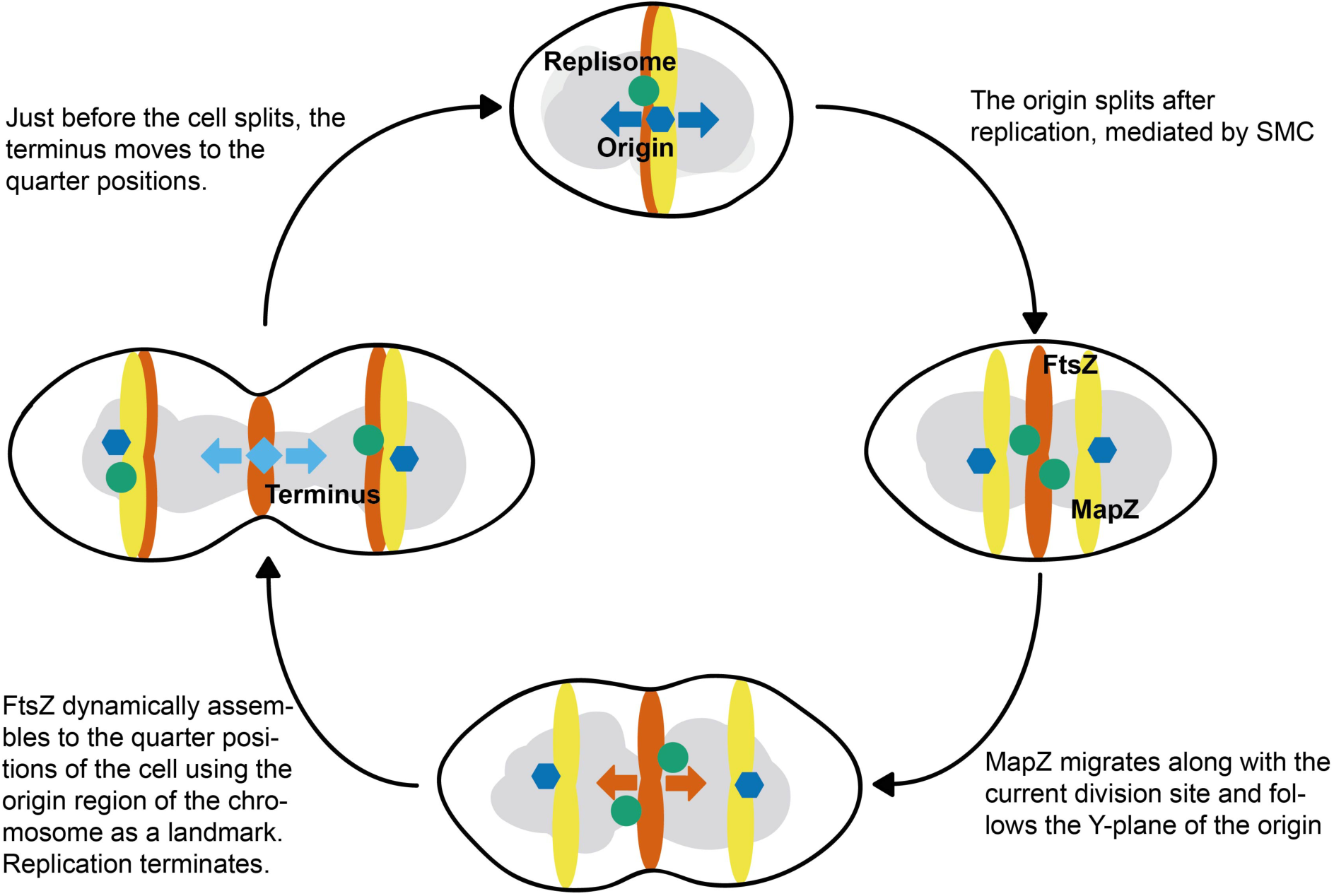
A schematic model for division site selection in pneumococci.

Recently it was found that MapZ localizes to future division sites before FtsZ and positions the Z-ring correctly via protein-protein interactions (*19*, *20*). We found that MapZ gradually moves with a similar timing as the chromosomal origins, but MapZ is not important for correct *oriC* positioning. On the other hand, perturbation of *oriC* segregation clearly resulted in altered MapZ localization, thus indicating the pivotal role of chromosomal origin positioning for proper cell division coordination. Notably, MapZ plays an important role in establishing the correct division plane, since the Z-ring frequently became non-perpendicular to the cell length axis when *mapZ* was deleted (Fig. 1d). Taken together, this suggests that while timely *oriC* positioning determines the timing of assembly and position of the cell division machinery, MapZ is a ruler for the correct angle of the division ring across the cell (Fig. 6). This explains the highly variable size and shape of *ΔmapZ* cells, as well as the cell division defect resulting from the mislocalized origins in Δ*smc* and ciprofloxacin-treated cells. Note, however, that although *oriC* segregation is clearly delayed in both the Δ*smc* strain as well as in the ciprofloxacin-treated cells, it eventually segregates to the future division sites in time before Z-ring assembly. This means that there are additional cues, and not solely SMC or topoisomerases which are involved in segregation and localization of *oriC,* explaining why the cell division defects resulting from *smc* deletion or ciprofloxacin treatment are not too severe. How the origin finds or simply remains attached to the future division site is unclear. We cannot rule out an as-of-yet unknown protein factor playing a role in this, for instance in keeping the newly replicated origins near the future division site. Perhaps coupled transcription-translation-transertion of membrane proteins encoded near *oriC* aid in transitory attachment of the chromosome to the membrane (*51*, *52*). Alternatively, physical, entropy-driven processes might be at play. In this respect, it is tempting to speculate that the origin region, which was recently shown to be highly structured and globular in shape (*53*), is pushed outside the region of active DNA replication and remains rather stationary in the crowded cytoplasm (*54*). The large globular structure of the origin can then act as a landmark for FtsZ polymerization and Z-ring formation. This hypothesis is in line with previous cytological observations demonstrating the absence of nucleoid occlusion in *S. pneumoniae* and efficient Z-ring formation over the nucleoid (*25*, *55*). The here-described division site selection mechanism by chromosomal organization is a simple way to coordinate DNA replication, chromosome segregation and division without the need for specialized regulators of FtsZ. Future research should reveal if this mechanism is also in place in other bacteria and whether the intimate relation between chromosome segregation and cell division can be used to treat bacterial infections using combination therapy targeting both processes.

## Acknowledgments

We thank Jeroen Siebring for initial work on *parB_p_*, Lieke van Gijtenbeek for providing *m(sf)gfp* and Oliver Gericke and Katrin Beilharz for technical assistance. We thank GeneCore, EMBL, Heidelberg for sequencing and Jelle Slager and Rieza Aprianto for help with analysis. Sequencing data is available at https://seek.sysmo-db.org/investigations/93#datafiles. We thank Dirk-Jan Scheffers for stimulating discussions and Sophie Martin and Stephan Gruber for constructive feedback on our manuscript. Work in the Veening lab is supported by the EMBO Young Investigator Program, a VIDI fellowship (864.12.001) and ERC starting grant 337399-PneumoCell. MK is supported by a grant from The Research Council of Norway (250976/F20).

## Supplementary materials

Materials and Methods

Figures S1-S9

Tables S1-S2

Movies S1-S3

References

## References

1. B. Alberts et al., in Molecular Biology of the Cell. (Garland Science, Taylor & Francis Group, New York, ed. 6th, 2014), pp. 963–1020.

2. J. P. Fededa, D. W. Gerlich, Molecular control of animal cell cytokinesis. Nat. Cell Biol. 14, 440–7 (2012).

3. A. H. Willet, N. A. McDonald, K. L. Gould, Regulation of contractile ring formation and septation in *Schizosaccharomyces pombe*. Curr. Opin. Microbiol. 28, 46–52 (2015).

4. J.-Y. Bouet, M. Stouf, E. Lebailly, F. Cornet, Mechanisms for chromosome segregation. Curr. Opin. Microbiol. 22, 60–5 (2014).

5. X. Wang, D. Z. Rudner, Spatial organization of bacterial chromosomes. Curr. Opin. Microbiol. 22, 66–72 (2014).

6. W. Margolin, FtsZ and the division of prokaryotic cells and organelles. Nat. Rev. Mol. Cell Biol. 6, 862–871 (2005).

7. V. W. Rowlett, W. Margolin, The Min system and other nucleoid-independent regulators of Z ring positioning. Front. Microbiol. 6, 478 (2015).

8. D. W. Adams, L. J. Wu, J. Errington, Cell cycle regulation by the bacterial nucleoid. Curr. Opin. Microbiol. 22, 94–101 (2014).

9. M. Thanbichler, L. Shapiro, MipZ, a spatial regulator coordinating chromosome segregation with cell division in *Caulobacter*. Cell. 126, 147–162 (2006).

10. J. Willemse, J. W. Borst, E. De Waal, T. Bisseling, G. P. Van Wezel, Positive control of cell division: FtsZ is recruited by SsgB during sporulation of *Streptomyces*. Genes Dev. 25, 89–99 (2011).

11. A. Treuner-Lange et al., PomZ, a ParA-like protein, regulates Z-ring formation and cell division in *Myxococcus xanthus*. Mol. Microbiol. 87, 235–253 (2013).

12. J. Männik, M. W. Bailey, Spatial coordination between chromosomes and cell division proteins in *Escherichia coli*. Front. Microbiol. 6, 306 (2015).

13. L. G. Monahan, A. T. F. Liew, A. L. Bottomley, E. J. Harry, Division site positioning in bacteria: one size does not fit all. Front. Microbiol. 5, 19 (2014).

14. A. Zaritsky, C. L. Woldringh, Chromosome replication, cell growth, division and shape: a personal perspective. Front. Microbiol. 6, 756 (2015).

15. I. V. Hajduk, C. D. A. Rodrigues, E. J. Harry, Connecting the dots of the bacterial cell cycle: Coordinating chromosome replication and segregation with cell division. Semin. Cell Dev. Biol. 53, 2–9 (2016).

16. M. Wallden et al., The synchronization of replication and division cycles in individual *E. coli* cells. Cell. 166, 729–739 (2016).

17. M. G. Pinho, M. Kjos, J.-W. Veening, How to get (a)round: mechanisms controlling growth and division of coccoid bacteria. Nat. Rev. Microbiol. 11, 601–14 (2013).

18. D. Fadda et al., *Streptococcus pneumoniae* DivIVA: Localization and interactions in a MinCD-free context. J. Bacteriol. 189, 1288–1298 (2007).

19. A. Fleurie et al., MapZ marks the division sites and positions FtsZ rings in *Streptococcus pneumoniae*. Nature. 516, 259–262 (2014).

20. N. Holečková et al., LocZ Is a new cell division protein involved in proper septum placement in *Streptococcus pneumoniae*. MBio. 6, e01700–14 (2014).

21. K. Beilharz et al., Control of cell division in *Streptococcus pneumoniae* by the conserved Ser/Thr protein kinase StkP. Proc. Natl. Acad. Sci. 109, E905–E913 (2012).

22. M. Bramkamp, Following the equator: division site selection in *Streptococcus pneumoniae*. Trends Microbiol. 23, 121–122 (2015).

23. C. Grangeasse, Rewiring the pneumococcal cell cycle with serine/threonine- and tyrosine-kinases. Trends Microbiol. (2016).

24. A. Minnen, L. Attaiech, M. Thon, S. Gruber, J.-W. Veening, SMC is recruited to *oriC* by ParB and promotes chromosome segregation in *Streptococcus pneumoniae*. Mol. Microbiol. 81, 676–688 (2011).

25. M. Kjos, J. W. Veening, Tracking of chromosome dynamics in live *Streptococcus pneumoniae* reveals that transcription promotes chromosome segregation. Mol. Microbiol. 91, 1088–1105 (2014).

26. R. A. Britton, D. C. Lin, A. D. Grossman, Characterization of a prokaryotic SMC protein involved in chromosome partitioning. Genes Dev. 12, 1254–9 (1998).

27. M. J. Boersma et al., Minimal peptidoglycan (PG) turnover in wild-type and PG hydrolase and cell division mutants of *Streptococcus pneumoniae* D39 growing planktonically and in host-relevant biofilms. J. Bacteriol. 197, 3472–3485 (2015).

28. A. Paintdakhi et al., Oufti: an integrated software package for high-accuracy, high-throughput quantitative microscopy analysis. Mol. Microbiol. 99, 767–777 (2016).

29. K. Beilharz, R. van Raaphorst, M. Kjos, J. W. Veening, Red fluorescent proteins for gene expression and protein localization studies in *Streptococcus pneumoniae* and efficient transformation with DNA assembled via the Gibson assembly method. Appl. Environ. Microbiol. 81, 7244–7252 (2015).

30. R. A. Daniel, J. Errington, Control of cell morphogenesis in bacteria: two distinct ways to make a rod-shaped cell. Cell. 113, 767–776 (2003).

31. L.-P. Potluri, M. A. de Pedro, K. D. Young, *Escherichia coli* low-molecular-weight penicillin-binding proteins help orient septal FtsZ, and their absence leads to asymmetric cell division and branching. Mol. Microbiol. 84, 203–224 (2012).

32. D. L. Popham, K. D. Young, Role of penicillin-binding proteins in bacterial cell morphogenesis. Curr. Opin. Microbiol. 6, 594–599 (2003).

33. S. Manuse et al., Structure-function analysis of the extracellular domain of the pneumococcal cell division site positioning protein MapZ. Nat. Commun. 7, 12071 (2016).

34. A. Fleurie et al., Interplay of the serine/threonine-kinase StkP and the paralogs DivIVA and GpsB in pneumococcal cell elongation and division. PLoS Genet. 10, e1004275 (2014).

35. K. P. Lemon, A. D. Grossman, Movement of replicating DNA through a stationary replisome. Mol. Cell. 6, 1321–1330 (2000).

36. R. Reyes-Lamothe, C. Possoz, O. Danilova, D. J. Sherratt, Independent positioning and action of *Escherichia coli* replisomes in live cells. Cell. 133, 90–102 (2008).

37. K. Jaqaman et al., Robust single-particle tracking in live-cell time-lapse sequences. Nat. Methods. 5, 695–702 (2008).

38. M. B. Elowitz, M. G. Surette, P. E. Wolf, J. B. Stock, S. Leibler, Protein mobility in the cytoplasm of *Escherichia coli*. J. Bacteriol. 181, 197–203 (1999).

39. U. Wegmann, K. Overweg, S. Jeanson, M. Gasson, C. Shearman, Molecular characterization and structural instability of the industrially important composite metabolic plasmid pLP712. Microbiology. 158, 2936–2945 (2012).

40. Y. Li, K. Sergueev, S. Austin, The segregation of the *Escherichia coli* origin and terminus of replication. Mol. Microbiol. 46, 985–996 (2002).

41. B. E. Funnell, L. Gagnier, P1 plasmid partition: binding of P1 ParB protein and *Escherichia coli* integration host factor to altered *parS* sites. Biochimie. 76, 924–932 (1994).

42. X. Wang, P. Montero Llopis, D. Z. Rudner, *Bacillus subtilis* chromosome organization oscillates between two distinct patterns. Proc. Natl. Acad. Sci. U. S. A. 111, 12877–12882 (2014).

43. P. H. Viollier et al., Rapid and sequential movement of individual chromosomal loci to specific subcellular locations during bacterial DNA replication. Proc. Natl. Acad. Sci. U. S. A. 101, 9257–9262 (2004).

44. B. Youngren, H. J. Nielsen, S. Jun, S. Austin, The multifork *Escherichia coli* chromosome is a self-duplicating and self-segregating thermodynamic ring polymer. Genes Dev. 28, 71–84 (2014).

45. S. Gruber, Multilayer chromosome organization through DNA bending, bridging and extrusion. Curr. Opin. Microbiol. 22, 102–110 (2014).

46. X. Wang et al., The SMC condensin complex is required for origin segregation in *Bacillus subtilis*. Curr. Biol. 24, 287–292 (2014).

47. S. Gruber et al., Interlinked sister chromosomes arise in the absence of condensin during fast replication in *B. subtilis*. Curr. Biol. CB. 24, 293–298 (2014).

48. E. Fernandez-Moreira, D. Balas, I. Gonzalez, A. G. de la Campa, Fluoroquinolones inhibit preferentially *Streptococcus pneumoniae* DNA topoisomerase IV than DNA gyrase native proteins. Microb. Drug Resist. 6, 259–67 (2000).

49. J. Slager, M. Kjos, L. Attaiech, J. W. Veening, Antibiotic-induced replication stress triggers bacterial competence by increasing gene dosage near the origin. Cell. 157, 395–406 (2014).

50. X. Wang, X. Liu, C. Possoz, D. J. Sherratt, The two *Escherichia coli* chromosome arms locate to separate cell halves. Genes Dev. 20, 1727–31 (2006).

51. C. L. Woldringh, The role of co-transcriptional translation and protein translocation (transertion) in bacterial chromosome segregation. Mol. Microbiol. 45, 17–29 (2002).

52. E. A. Libby, M. Roggiani, M. Goulian, Membrane protein expression triggers chromosomal locus repositioning in bacteria. Proc. Natl. Acad. Sci. U. S. A. 109, 7445–50 (2012).

53. M. Marbouty et al., Condensin- and replication-mediated bacterial chromosome folding and origin condensation revealed by Hi-C and super-resolution imaging. Mol. Cell. 59, 588–602 (2015).

54. B. R. Parry et al., The bacterial cytoplasm has glass-like properties and is fluidized by metabolic activity. Cell. 156, 183–194 (2014).

55. A. D. Land et al., Requirement of essential Pbp2x and GpsB for septal ring closure in Streptococcus pneumoniae D39. Mol. Microbiol. 90, 939–955 (2013).

